# Ribozyme synthesis of both L- and D- amino acid oligos

**DOI:** 10.1101/2023.04.28.538729

**Authors:** Yuhong Wang

## Abstract

The ribosome is responsible for assembling proteins using 20 naturally occurring L-handed amino acids. However, incorporating non-natural amino acids into a protein is a challenging process needs improvement. In this study, we report a new possible approach to creating nonnatural peptides using ribozymes inspired by the peptidyl transfer center. These RNA scaffolds, which are approximately 100 nucleotides in length, bind to RNase T1 truncated tRNA-like chimeras and bring them into close proximity to facilitate peptide ligation. We used single-molecule fluorescence resonance energy transfer (smFRET) to show close distances between RNA-RNA, tRNA^Lys^-tRNA^Lys^, and RNA-tRNA^Lys^ pairs, which strongly suggests that the mechanism of peptide ligation is due to the proximity of the substrate through dimerization of the enzymes. Mass spectrometry analysis confirmed the detection of oligopeptides from four amino acids, including L-Lysine, D-Lysine, L-Phenylalanine, and D-Phenylalanine. These results indicate that ribozymes have greater flexibility in accommodating nonnatural amino acids. Our findings pave the way for potentially new avenues in the synthesis of nonnatural peptides, beyond the limitations of ribosomal peptide synthesis and other existing methods.

**GRAPHICAL ABSTRACT:** 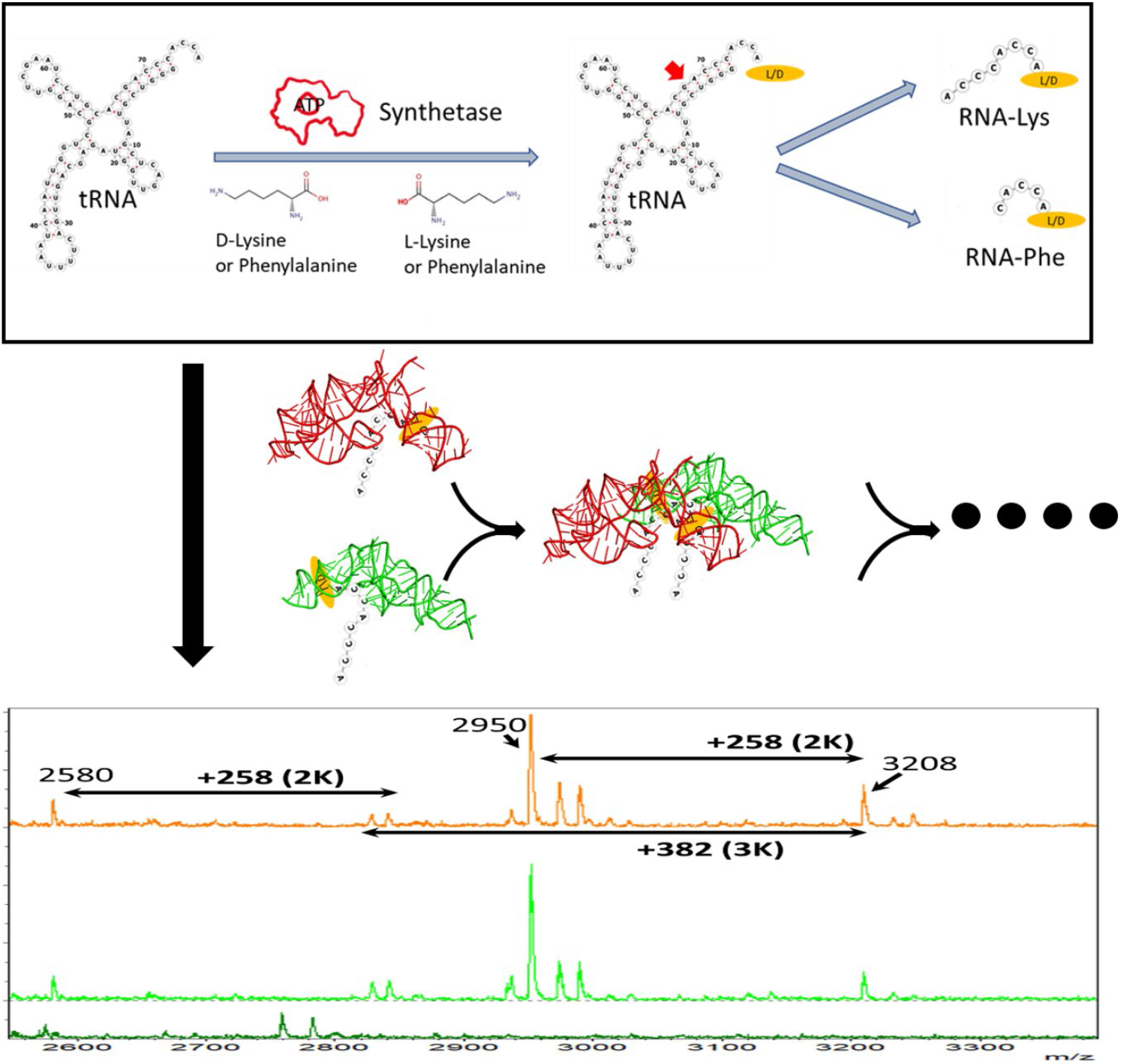

## INTRODUCTION

Ribosome is responsible to make proteins in all three-domain cells. However, the ribosome itself contains proteins, making it difficult to extrapolate the protoribosome prior to the protein world. This remains challenging because the ribosome is mostly developed at the time of the last common ancestor, LUCA.^1^ The peptidyl transfer center of the ribosome (PTC) is highly conservative and most ancient based on comparative sequence analysis.^2^ The finding of a 2-fold rotational symmetry between the tRNA A- and P-binding site at the PTC implies gene duplication as the origin of ancient ribosome.^3^ However, these structurally symmetric parts, named ptc1a and ptc1b (Figure 1C) are not superimposable on their primary sequence.^4^ As shown in Figure 1b, large insertions are developed in the ptc1a sequence, making the experimental demonstration of the dimerization and gene duplication hypothesis elusive. Recently, we have identified two RNA scaffolds that have the ability to dimerize and connect oligopeptides.^5,6^ Unlike previous attempts to create symmetrical sequences from a 2D map, we discovered a superimposable sequence by aligning the A- and P-site tRNA while maintaining the ribosomal RNAs as part of the objects with the tRNAs.^4^ Upon superimposing the tRNA parts, the ptc1a and ptc1b parts were also superimposed. The insertions in the ptc1a were then trimmed using the superimposed ptc1b as the guide. Finally, the ptc1a and ptc1b were superimposed again without other parts and the gaps due to trimming in ptc1a were filled with sequence from ptc1b. This is done with pymol’s cealign command using ribosome structure 4wpo (Figure S1).^7,8^ The final sequences are displayed in Figure 1A. Using labeled ptc1a/b and tRNA-like chimeras, we have observed high smFRET signals to indicate close proximity between ptc1a/1b,the chimeras, and between ptc 1b and tRNA chimeras, which led us to explore the possibility of peptide formation.^5^ Strikingly, we detected peptide oligomers using MALDI-TOF Mass Spec, suggesting that ribozymes may serve as an alternative way of synthesizing peptide chains. This discovery has important implications for both biotechnology and evolution research, as it could lead to new methods for synthesizing peptides and proteins and shed light on the early stages of protein evolution and the origins of life. Here, we report the formation of oligopeptides from nonnatural amino acids.

**Figure 1.**
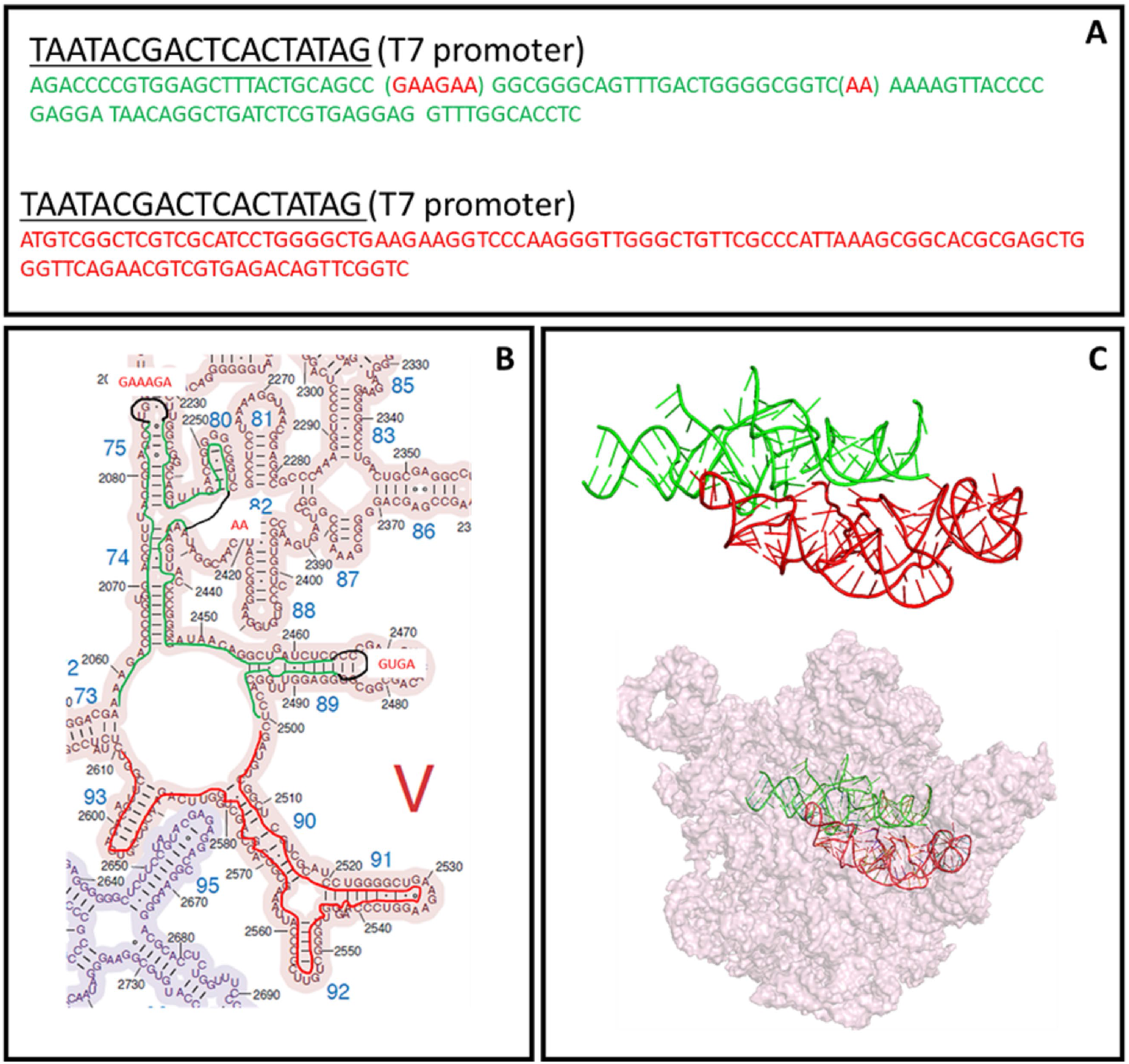
(A) sequences of the ribozymes in the main text. The top and bottom are sequences for ptc1a and ptc1b, respectively. (B) Secondary structure of ptc1a and 1b and their positions on the 2D map. The modifications are shown with black lines and red-font residues. (C) the 3D structure of the ptc1a/1b and their positions at the peptidyl transfer center.

A variety of approaches have been employing to modify ribosome system to incorporate nonnatural amino acid, including the introduction of orthogonal tRNA-synthetase pairs, reassignment of degenerate codons, and modification of the peptidyl transfer center. Despite these advances, significant obstacles remain, such as low efficiency and stalling during translation.^9-13^ On the other hand, nonribosomal peptide synthetases (NRPS) are capable of synthesizing a wide range of peptide antibiotics and other biologically active compounds. These enzymes are large and modular, working in linear arrays, which makes module shuffling, splitting, and insertion challenging.^14,15^ On the other hand, the ribozymes that we report here may have the potential to offer a complementary approach to the existing methods.

## MATERIAL AND METHODS

All the DNA templates for in vitro transcriptions of ptc1a/1b, are purchased from IDTDNA. Lysine and phenylalanine specific tRNAs are purchased from chemical-blocks.com (obsoleted). NHS dyes are purchased from lumiprobe. Radioactive amino acids are purchased from PerkinElmer. Other reagents are from Millipore Sigma if not specified. The T7 transcription, RNA labeling and biotinylation, tRNA synthetase expression, tRNA charging, Rnase T1 digestion and gel purification are described in the supporting information.

### Ribozyme reaction and MALDI-TOF mass spec

The ribozyme reactions are mixtures of 1-10 microM of ptc1a/b, 10-20 mM MgCl_2_, 2-10 A_260_/ml RNA-AA chimeras. The reactions are incubated at 37 °C for various time intervals (t=0, 2hr), and the products are analyzed by Bruker autoflex speed MALDI-TOF mass spectroscopy.

To prepare for mass spec, the samples were bound to ziptip (Agilent) and washed with 0.1M Triethylamine acetate and water. The samples are spotted on MTP standard 384 plate together with saturated Sinapic acid in 25 mM ammonium citrate and 70% Acetonitrile. The method is reflectron negative mode. Every set of data is calibrated by commercial standard RNA spotted multiple locations on the same plate.

### smFRET measurement

The fluorescent microscope is modified on a Nikon eclipse Ti2-E inverted microscope with two CMOS cameras and perfect-focusing-system as previously described. ^5 16^ In this newer version microscope, there are two turrets. The top turret reflects the laser illumination toward the objective above the total internal reflection angle, which generates the evanescent excitation field. The fluorescence via FRET and direct excitation are collected by the same objective, passed through the first turret, and split by the second turret; and eventually reached the two cameras. Because these fluorescence signals are from same molecule but split spectroscopically, the physical positions on the cameras are strictly correlated with a single pair of offset values (delta-X and delta-Y) for all the FRET pairs. The FRET pairs’ fluorescence intensities are fitted with Nikon Elements program and FRET value is calculated as I_acceptor_/(I_acceptor_ + I_donor_). FRET values larger than 50% indicate distance closer than R0. For Cy3/Cy5 pair, this R0 is approximately 5 nm. Typically, about 40 fields of view are collected. In each field, about 1,000 FRET pairs are observed in 10s with 100 ms intervals. For all measurements, samples are diluted to the range of 10-100 nm. An oxygen scavenger cocktail is added to the channels before imaging to prevent dye photobleaching.

## RESULTS

### Synthesis of RNA-AA substrates

Our previous work showed that while ribozyme dimerization can mimic the function of ribosomes by bringing the substrates together, intact tRNAs are not suitable for this process. Only short RNA-AA chimeras are accepted (AA stands for amino acid). To address this, the first step is to prepare these short RNA-AA chimeras. In addition to natural lysine and phenylalanine, we selected two non-natural amino acids, D-lysine, and D-phenylalanine, as L- and D-forms are believed to be indistinguishable in protoribosomes.^1^

Figures 2A depicts the preparation scheme for RNA-lysine and RNA-phenylalanine chimeras. The purified lysine or phenylalanine synthetase is mixed with either D- or L-lysine or phenylalanine, respectively, along with their corresponding tRNAs. Previous research has shown that natural tRNA synthetases can charge many non-natural amino acids.^17-20^ However, charging of D-lysine/phenylalanine has only been demonstrated through the flexizyme method and chemical synthesis. ^20,21^ Our experiments indicate that these D-amino acids are charged by their L-amino acid synthetase to a reasonable extent. The RNA-AA chimeras are subsequently digested with RNase T1 at the “G” residues (Figure 2A). For lysine-tRNA^Lysine^ this generates ACCCACCA-Lysine, while it produces CACCA-phenylalanine for Phenylalanine-tRNA^phenylalanine^. These RNA-AA chimeras are not precipitable and are therefore purified through gel separation, as shown in Figure 2B. The T1 digestion mixture of ^14^C-phenylalanine charged tRNA generates three bands that migrate faster than bromophenol blue; and two bands between this dye and xylene cyanol FF. In lane one, a commercial CACCA standard migrates at the same position as the lower band between bromophenol blue and xylene cyanol FF. And this band also showed radioactivity after band-incision and elution. The MALDI-TOF Mass spec displays the expected mass peak at 1660 Da (2C). Likewise, we have identified the RNA-AA with lysine. Representative preparative mini gels (1.5 mm) are shown in Figure S2. Subsequently, we prepared the non-radioactive RNA-AA chimeras, including non-radioactive ^13^C and ^15^N labeled phenylalanine and lysine, respectively. The species of L-, D-, and ^13^C-L-phenylalanine exhibit 1660, 1660, 1661 Da peaks in Figure 2D as orange, green, and blue traces, respectively. Likewise, the species of L-, D-, and ^15^N, ^15^N-L-phenylalanine exhibit 2580, 2580, 2582 Da peaks in Figure 2E as orange, green, and blue traces, respectively. For RNA-Phe(L), additional tandem MS/MS spectra were taken, which is displayed in Figure S3. The isotopic shifting is strong evidence to support the assignment of the proper chemical species.

**Figure 2.**
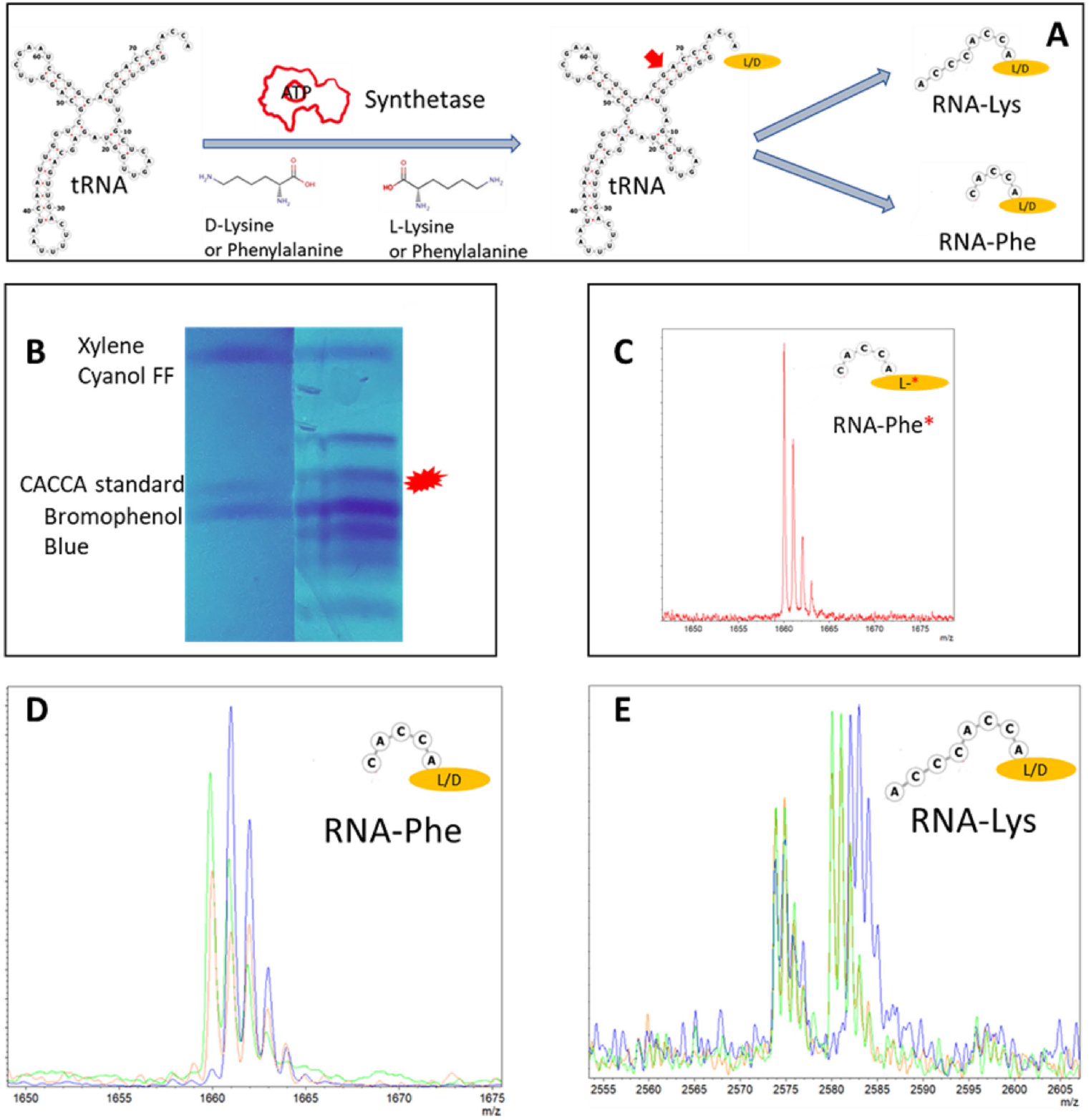
(A) Natural synthetases of lysine RS and phenylalanine RS are able to charge their natural substrates and the D-analogs. (B) 20% PAGE gel to purify the charged, RNase T1 digested RNA-AA chimeras. Lane 1: loading dyes and CACCA standard. Lane 2: Migration bands after RNase T1 digestion of tRNA-phenylanaline (^14^C labeled.) The band that matches the standard migration position exhibited radioactivity. (C) MALDI-TOF Mass spectrum of the eluted RNA-AA from the radioactive band in (B). The read * indicates 14C radioactivity. (D) MALDI-TOF Mass spectrum of charged CACCA-Phenylalanine. The orange, green and blue spectrum are for L-, D-phenylalanine, and L-13C-labeled phenylalanine, respectively. (E) MALDI-TOF Mass spectrum of charged ACCACCA-lysine. The orange, green and blue spectrum are for L-, D-lysine, and L-15N,15N-labeled lysine, respectively.

Figure 2 shows that we have successfully generated short RNA-AA chimeras through enzymatic means. This type of chimera is challenging to prepare, but it is a necessary prerequisite for ribozyme ligation. Other methods, such as chemical modifications and flexizymes, have been developed, but they require extensive synthetic expertise or activated substrates.^22,23^ However, we took advantage of natural synthetases, which does not require additional activation or modification steps. The charging efficiencies of the D-amino acids were not directly measured due to the lack of radioactive amino acids, but several parallel mass spectrometry experiments were performed, and the signals were comparable. For example, Figure S4 showed the percentage of charged RNA vs. sum of charged and uncharged RNA are approximately 30-50% for D- and L-AAs. This percentage agrees from the radioactivity estimate of full length tRNA charging.

### Synthesis of Peptide Oligos

After preparing and confirming the substrates, they were mixed with ptc1a/b and MgCl_2_ at 37ºC for 2 hours. The formation of oligos was monitored using mass spectrometry. Figures 3A shows examples of poly-lysine formation. We observed RNA-Lysine peak at 2580 Da, and higher mass peak shifted 2 lysine molecular weight (MW, 2837). Similarly, there is another mass-shift of 2 lysine between 2948 Da and 3205 Da peaks. However, the difference between 2948 Da to 2837 Da is 17 Da less than the expected MW of 1 lysine adduct, which could be due to loss of “NH_2_” on a lysine side chain. Both L- and D-lysine exhibited similar mass spectra. In addition, given the potential for RNA fragmentation during MALDI-TOF conditions, RNA-(lysine)n on shorter RNAs were searched for and are depicted in Figure S5. In this figure, oligopeptides with n=1, 2, 4, 5 were observed attached to CCACCA. Figures 3B shows the similar mass detection for poly L- and D-phenylalanine. There are good agreements between the theoretical prediction and experimental results for RNA-(phenylalanine)_n=1,2,3,5_. In addition, ptc1a/b only control is presented in Figure 3B as well (the bottom trace).

**Figure 3.**
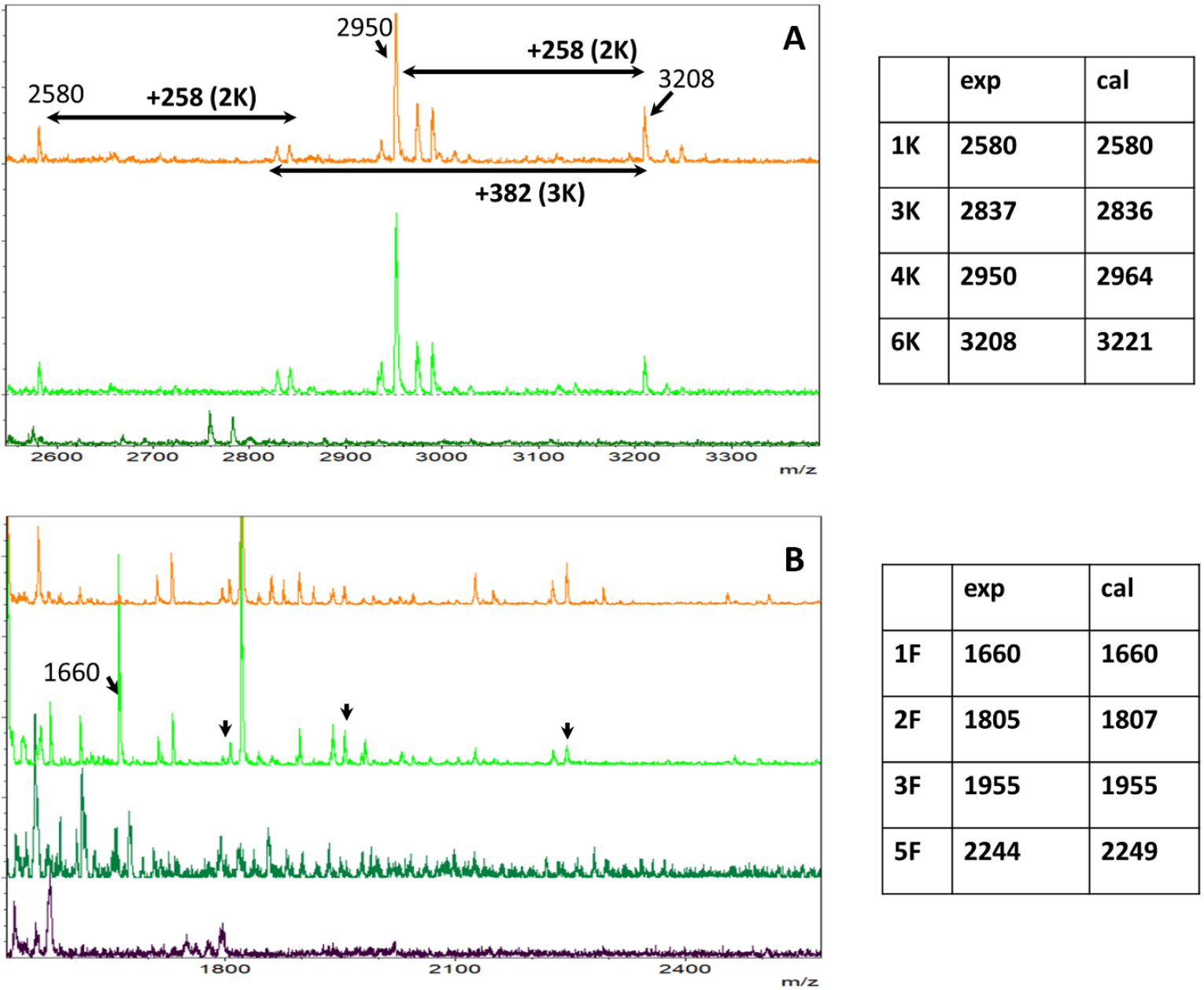
(A) MALDI-TOF spectrum of reaction of CACCACCA-(Lysine)_n_. The major peaks are highlighted and their experimental and calculated values are compared. For bottom to top, the traces are T=0 for L-lysine reaction, T=2hr for D-lysine reaction, and T=2 hr for L-lysine reaction. (B) MALDI-TOF spectrum of reaction of CACCA-(Phenylalanine)_n_. The major peaks are highlighted and their experimental and calculated values are compared. For bottom to top, the traces are T=2hr for ptc1a/b only control, T=0 for L-phenylalanine reaction, T=2hr for D-phenylalanine reaction, and T=2 hr for L-phenylalanine reaction.

The fact that the oligopeptide peaks appear only after incubation in Figures 3A and B, compared to time zero controls, eliminates any contribution from ptc1a/1b and confirms that the peptide oligos are indeed the product of the reaction. The ptc1a/b appear stable during 2-hour incubation time, because the ptc1a/b control trace in Figure 3B did not show any significant peaks either. In Figures 3A and 3B, as well as in Supplementary Figure S5, only RNA-peptides on authentic RNA fragments are highlighted. However, there are many more mass differences that match peptide oligos perfectly, but do not match the intact RNA-AAn formula. This is due to the complex fragmentation of RNA during MALDI-TOF.

### smFRET study to reveal substrate proximity mechanism

The mechanism of substrate proximity during the formation of oligopeptides by the ribozyme was observed using fluorescence resonance energy transfer (FRET) with Cy3/Cy5 dyes labeled lysine side chains (Figure 4). Similar to unlabeled RNA-lys (Figure S2), Cy3/Cy5-labeled RNA-lys bands are incised and eluted. The subsequent mass measurement confirms the species. It is used for the smFRET experiments. The experiments were conducted using a total internal reflection fluorescence microscope, based on a Nikon Ti2E with dual turrets and cameras (Figure 5A). The ribozymes were tethered on a biotinylated surface via biotin-streptavidin-biotin interaction, and interactions between ribozyme-substrate or substrate-substrate were indicated by the FRET signals. The fluorescence emissions from the donor and acceptor dyes were collected by different cameras, and the physically overlapped spots in the two camera images indicate the presence of a single ribozyme complex with both Cy3 and Cy5 labeling, similar to previous studies.^5^ In Figure 5B, one representative image is shown, in which the green and red dots are fluorescence from donor and acceptor, respectively. We have observed high FRET efficiencies around 0.54-0.55 (48.3-48.7 Å) between Cy3-L-lysine (or Cy3-D-lysine)-RNA and Cy5-(D-lysine)-RNA. In addition, FRET between 3’-Cy3 labeled ribozyme (named ptc1b) and Cy5-DK-ACCCAC is approximately 0.57 (approximately 47.4 Å).

**Figure 4.**
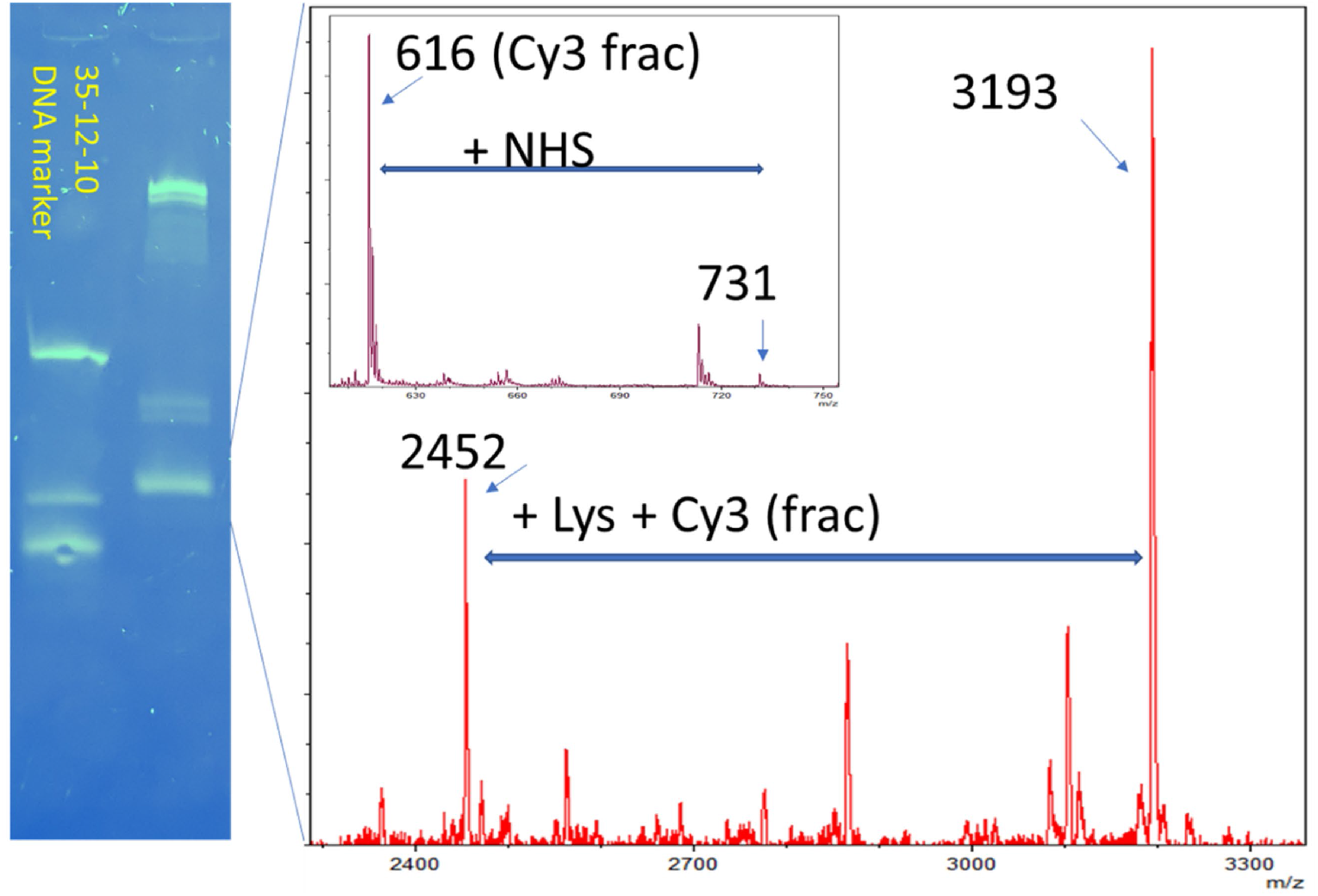
Gel separation of Cy3-labeled RNA-lys. The MALDI-TOF spectrum shows mass peak at 3193 (theory at 3196). The insert is the mass fragment of the Cy3-dye only. The mass at 731 Da is the intact dye and 616 Da is the dye after losing NHS group.

**Figure 5.**
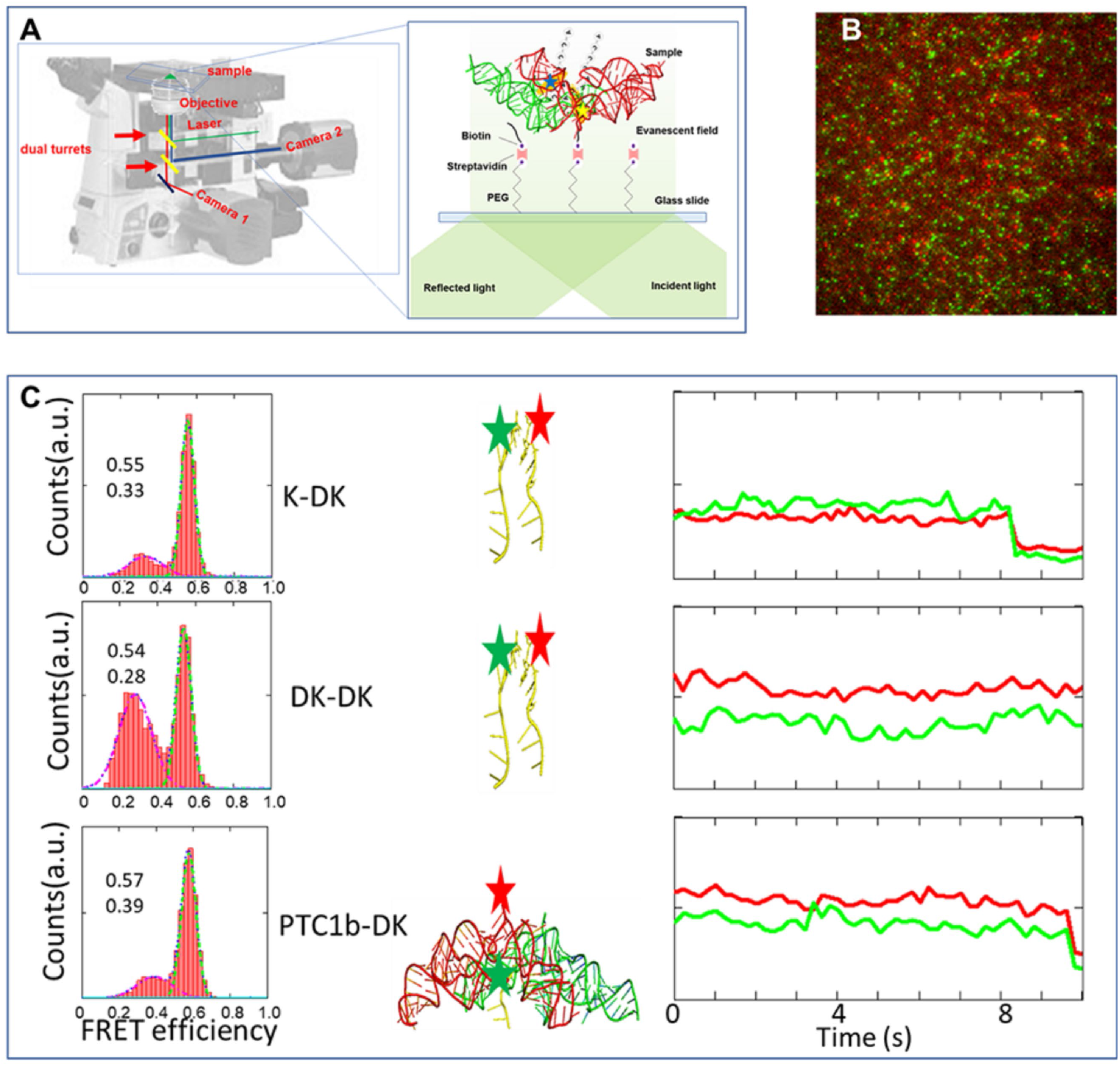
(A) TIRF based smFRET instruments. FRET-paired dyes are labeled on the ribozymes or their substrates. The biomolecules are tethered on the glass surface and observed one-by-one under laser illumination. (B) one representative overlay image. The red and green dots are fluorescence emissions from the sample biomolecule that are separated spectroscopically. (C) FRET efficiencies histograms between L-lysine (or D-lysine)-RNA and (D-lysine)-RNA, as well as between ribozyme ptc1b and (D-lysine)-RNA. These high FRET species indicate close substrate proximity inside the ribozyme. The red and green stars indicate the position of Cy5/Cy3 dye labeling, respectively. For each FRET experiments, one representative fitted time-lapse donor/acceptor fluorescent intensities are shown in green/red, respectively. FRET efficiency = *I*_acceptor_/(*I*_acceptor_+*I*_donor_).

Previously, we have published ∼ 0.6 FRET efficiencies in similar experiments for L-lysine chimeras. The results here show that the ribozyme can tolerate D-lysine, contrary to the strict selection of ribosomes. The high FRET signals observed between the RNA-AA and RNA-AA, as well as between RNA-AA and ptc1b, suggest that the mechanism of oligo formation is similar to that of the ribosome. There is a smaller peak ∼ 0.3 FRET efficiency (57.6 Å), suggesting some of these complexes are in different orientation or dimerize with larger intermolecular distances. As the dyes are tethered to the lysine moiety, the results are clear indicators of RNA-AA molecules, including non-natural amino acids.

Our previous studies, using FRET-pair dyes labeling at various locations, characterized the interaction between ptc1a/1b and alternative tRNA-like chimeras, such as RNase T1 digested fragment, minihelix, uncharged chimeras, as well as the competition for binding in the presence of chloramphenicol. We also found some of the DNA counterparts could interact similarly. By labelling the DNA/RNA molecules at central and peripheral positions and mixing them with three different types of tRNA-like oligos, we observed that both DNA and RNA scaffolds can bring together several short tRNA-acceptor-domain analogs but not full-length tRNAs to close proximity. Magnesium ions and continuing three-way junction motifs are essential to this process, but amino acid charging to the tRNA analogues is not required.^5^ Figure 5 directly compares L- and D-lysine labeled at their amino acid moieties. These comparisons are unambiguous because the labelings are only on the amino acids. Therefore, these smFRET data strongly support the high flexibility of the ribozyme towards non-natural amino acids.

## DISCUSSION

In this study, we have demonstrated the feasibility of synthesizing non-natural peptides without the restriction of ribosomes. This new ribozyme complements existing methods, such as the bioengineering of ribosomes and tRNAs, which aim to modify but not abolish ribosomal mechanisms, and non-ribosomal peptide synthases, which are still in the early stages of development. Additionally, chemical synthesis is costly, environmentally unfriendly, and has limited available amino acids. While ribozyme synthesis of non-natural peptides is not yet advanced, it has the potential to provide a new approach for expanding the scope of drug-leading peptides beyond the limitations of ribosomes and other methods.

The unique combination of the 3-way junction (H74-75 and H80 for ptc1a, H90-92 for ptc1b in Figure 1) and bi-functional tRNA mimics (acting as both donor and acceptor for amide bond formation) results in a highly specific ribozyme system that can efficiently synthesize oligopeptides. As shown in Figure 6, the A-site tRNA is positioned via the loop on Helix 92 in ptc1b, while Helices 90,91 and 93 form a specific coaxial structure. The two terminals H91 and H93 dimerize with another ptc molecule’s H93 and H91, or ptc1a’s equivalents, respectively. Hence, place two H92 loops close. Therefore, we propose that these are the minimal ribozyme elements to enable continuing ligation. On the other hand, it is obvious that bi-functional substrates are necessary to polymerize. Additionally, several ribozymes that contain stem-loop-stem structures but lack 3-way junction, were found making single bond between puromycin mimics and can also dimerize.^24^ it will be interesting to investigate whether these ribozymes with stem-loop-stem structures can form multiple peptide bonds when given bi-functional substrates. Nevertheless, these findings are consistent with our previous results on dimerization and reinforce the idea that these ribozymes are capable of polymerizing peptides. However, in their studies, dimerization and peptide bond formation were detected by Gel-shifting and by Mass Spec, separately. There was no direct evidence to connect these two results regarding mechanism. On the contrary, smFRET provides a direct link between the formation of the peptide bond and the dimerization of the ribozyme, offering strong evidence for the mechanism of peptide bond formation. The use of smFRET to monitor the interaction between the substrate, ribozyme, and antibiotic has allowed us to offer a more comprehensive understanding of the ribozyme’s behavior and its ability to synthesize nonnatural peptides. The observation that chloramphenicol, which binds at the peptidyl transfer center, disrupted the proximity between the substrates but not between the ribozymes themselves further supports this conclusion. This result is unique to our study and sets it apart from other literature in the field.^6^

**Figure 6.**
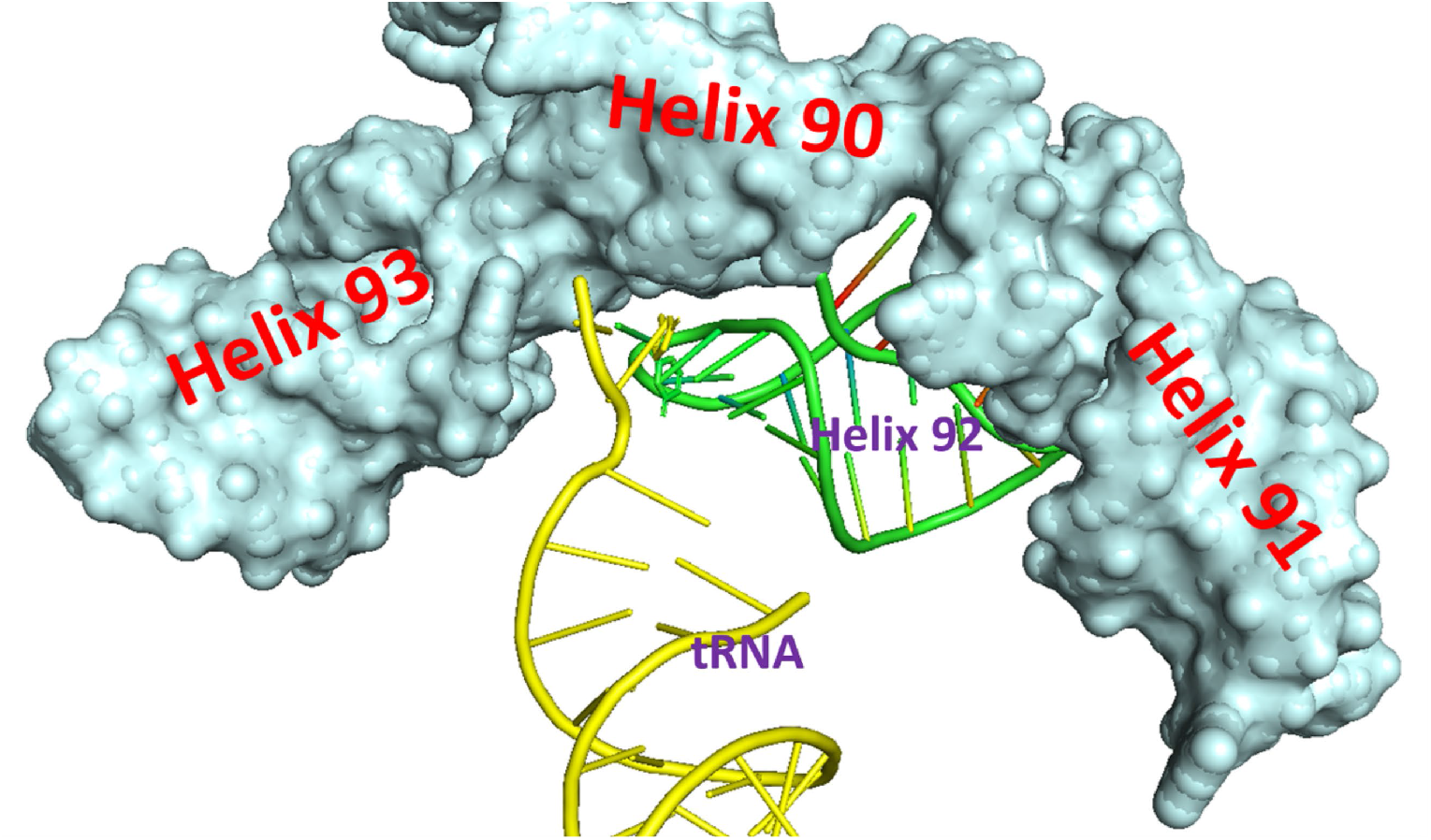
Coaxial structure of the helices of 90, 91 and 93 in ptc 1b enables the specific position of helix 92, which may be important to place the tRNA substrates at optimal reaction coordinates.

This report is an early step toward efficient ribozymes, which at the present has low yield of peptide formation. This caveat limits the method to detect the peptide other than Mass spec. We expect further improvements in ribozyme efficacy through in vitro evolution and saturation mutagenesis.^25,26^ The other crucial factor to this method is the synthesis of short RNA-AA chimeras. Although the use of RNase T1 digestion on natural or minihelix after charging provides acceptable substrates, direct charging of AA to shorter RNAs will eliminate unwanted RNA pieces from RNase T1 slicing, resulting in stronger mass spec signals and reducing ambiguity in peak assignment. Despite future challenges, our results present promising advancements in realizing enzymatic peptide synthesis without the limitations of ribosomes.

The discovery of protein-free peptide ligases has fundamental importance in understanding the evolution and origins of life. Simple ribozymes have already demonstrated the proof-of-concept, but the exploration of ribozymes capable of forming multiple peptide bonds has never been studied or considered for practical applications. Our reported ribozymes are capable of forming multiple peptide bonds due to their unique RNA sequences and the use of dual-functional substrates, whereas other studies utilized puromycin-like mimics. Additionally, in terms of the evolution of the ribosome, it is crucial to demonstrate that the proximity catalytic mechanism is shared by the ribosome and its precursor. To the best of our knowledge, our single-molecule fluorescence resonance energy transfer (smFRET) experiments are the only ones to have successfully achieved this goal.

Supplementary Data are available at NAR online.

## ACKNOWLEDGEMENTS

Y.W. thank Rice University’s “SEAS” program to use their Bruker autoflex MALDI-TOF instrument.

## DATA AVAILABILITY STATEMENT

Data available upon

