## supporting information for "Ribozyme synthesis of both L- and D- amino acid oligos"

Yuhong Wang[1] *

[1]Department of Biology and Biochemistry, University of Houston, Houston, TX 77204, USA

This supporting information contains details of material preparations and 5 supplemental figures, 1 supplemental tables.

Table of contents:


| Content | page |
| --- | --- |
| Title | 1 |
| Table of contents | 2 |
| Figure S1 | 3 |
| Figure S2 | 4 |
| Figure S3 | 5 |
| Figure S4 | 6 |
| Figure S5 | 7 |
| Experimental procedures | 8 |
| Reference | 11 |

Figure S1.

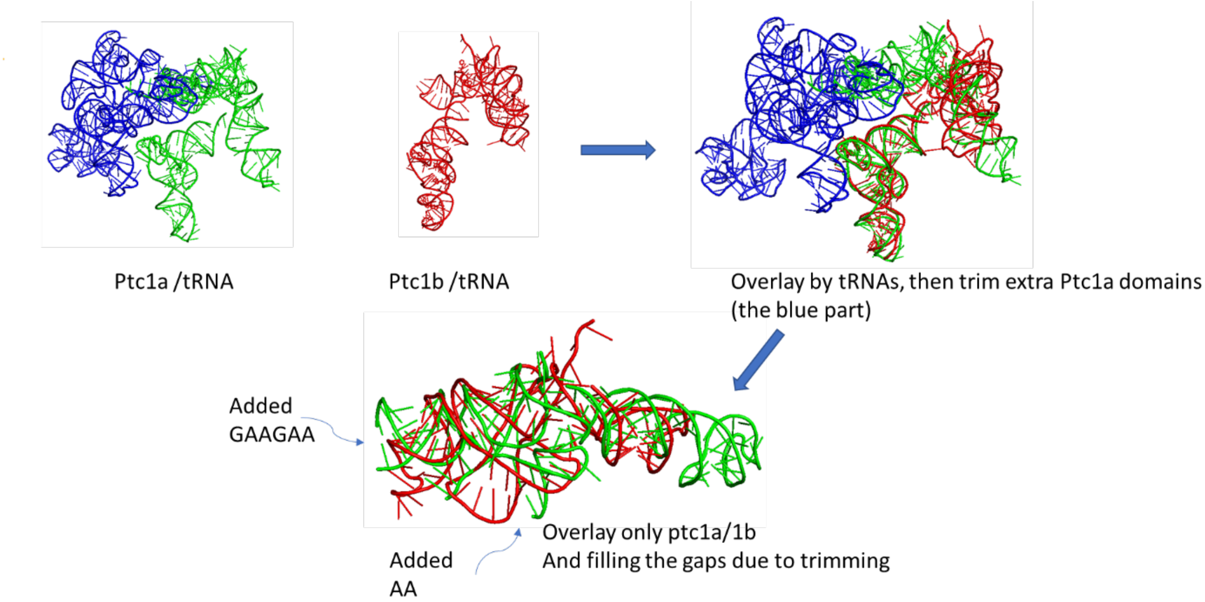


**Figure S1.** The process to identify ptc1a and ptc1b. These two RNAs were first included in two objects with tRNAs. While the tRNAs were aligned, the RNA parts coarsely aligned as well. Then the insertions of ptc1a were trimmed away guided by ptc1b. Finally, the two RNA pieces were re-aligned and the gaps in ptc1a were filled with similar residues in ptc1b.

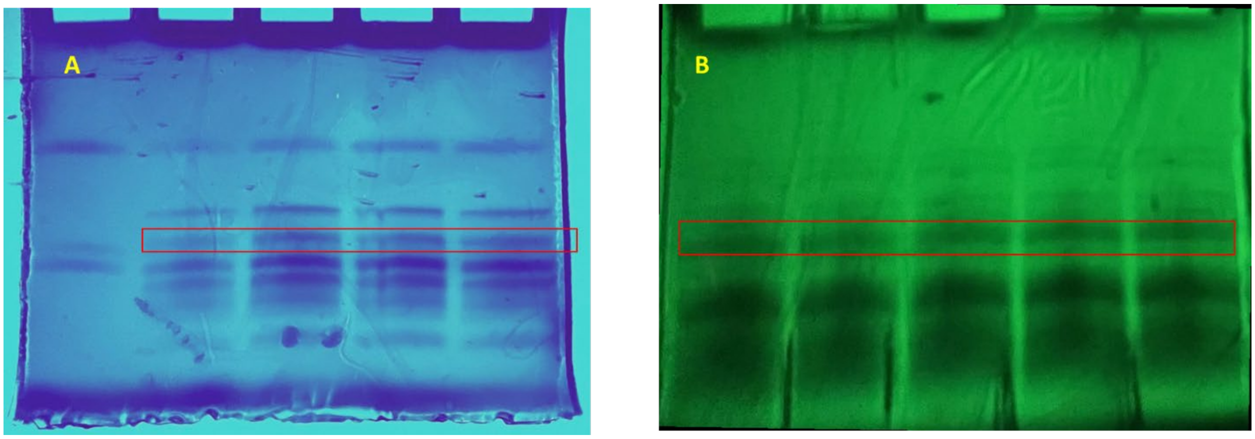


**Figure S2.** Preparative Gel images for RNase T1 digestion of (A) phenlalanine-tRNA$^{phenylalanine}$; (B) lysine-tRNA$^{lysine}$. The red boxes indicate the bands that are incised and eluted to obtain the RNA-AA chimeras. In experiment (B), no loading dyes are present. These images are obtained by 260 nm UV radiation of Gels placed on fluorescent silica TLC plate.

Figure S3.

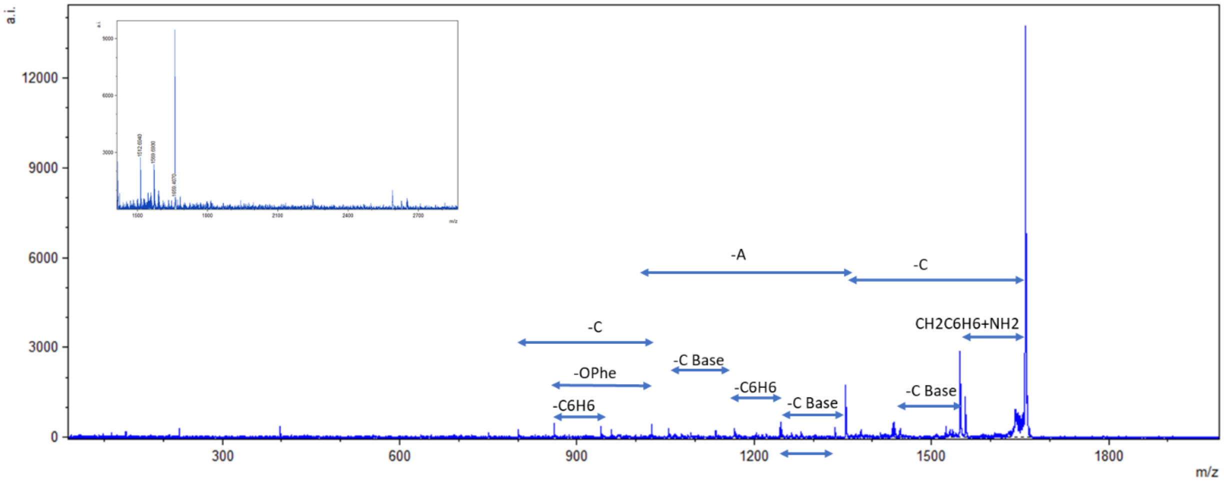


**Figure S3.** Mass and tandem MS/MS fragment of CACCA-Phenylalanine. The insert showed the major molecular peak at 1660 Da. And the arrows showed the assigned fragments of the MS/MS on this peak. The full mass list of the MS/MS is shown below:

| m/z | int. | r. int. | s/n | z | fwhm | resol. |
|---|---|---|---|---|---|---|
| 225.8390 | 283 | 2.54 | 14.7 | | 0.3922 | 576 |
| 397.0540 | 308 | 2.77 | 16.8 | | 0.3390 | 1171 |
| 860.3190 | 345 | 3.10 | 16.3 | | 0.4436 | 1939 |
| 940.2150 | 267 | 2.40 | 11.5 | | 0.4374 | 2150 |
| 1025.2900 | 351 | 3.15 | 13.9 | | 0.5095 | 2012 |
| 1054.1580 | 245 | 2.21 | 9.5 | | 0.6031 | 1748 |
| 1165.0710 | 224 | 2.02 | 8.2 | | 0.6107 | 1908 |
| 1242.9620 | 258 | 2.32 | 8.5 | | 0.5963 | 2084 |
| 1244.9350 | 409 | 3.68 | 13.5 | | 0.6748 | 1845 |
| 1245.9050 | 222 | 1.99 | 7.3 | | 2.4508 | 508 |
| 1335.8330 | 249 | 2.23 | 7.2 | | 0.6098 | 2191 |
| 1353.8210 | 1422 | 12.78 | 38.2 | | 0.6804 | 1990 |
| 1354.7980 | 848 | 7.62 | 22.7 | | 1.8068 | 750 |
| 1355.8480 | 301 | 2.70 | 8.0 | | 3.0795 | 440 |
| 1433.6160 | 345 | 3.10 | 7.4 | | 0.6310 | 2272 |
| 1434.6220 | 264 | 2.37 | 5.6 | | 0.6594 | 2176 |
| 1436.6360 | 411 | 3.69 | 8.7 | | 0.8330 | 1725 |
| 1547.2760 | 2356 | 21.16 | 29.7 | | 0.7658 | 2020 |
| 1548.3230 | 1487 | 13.36 | 18.6 | | 1.9537 | 792 |
| 1549.4090 | 548 | 4.92 | 6.8 | | 3.3492 | 463 |
| 1556.2420 | 1084 | 9.74 | 12.8 | | 0.7990 | 1948 |
| 1557.2530 | 610 | 5.48 | 7.2 | | 1.9661 | 792 |
| 1657.9750 | 11130 | 100.00 | 74.5 | | 0.8064 | 2056 |
| 1659.0190 | 7517 | 67.54 | 51.2 | | 2.0202 | 821 |
| 1660.1430 | 3173 | 28.51 | 22.0 | | 3.5459 | 468 |
| 1661.1480 | 1205 | 10.83 | 8.5 | | 5.0984 | 326 |

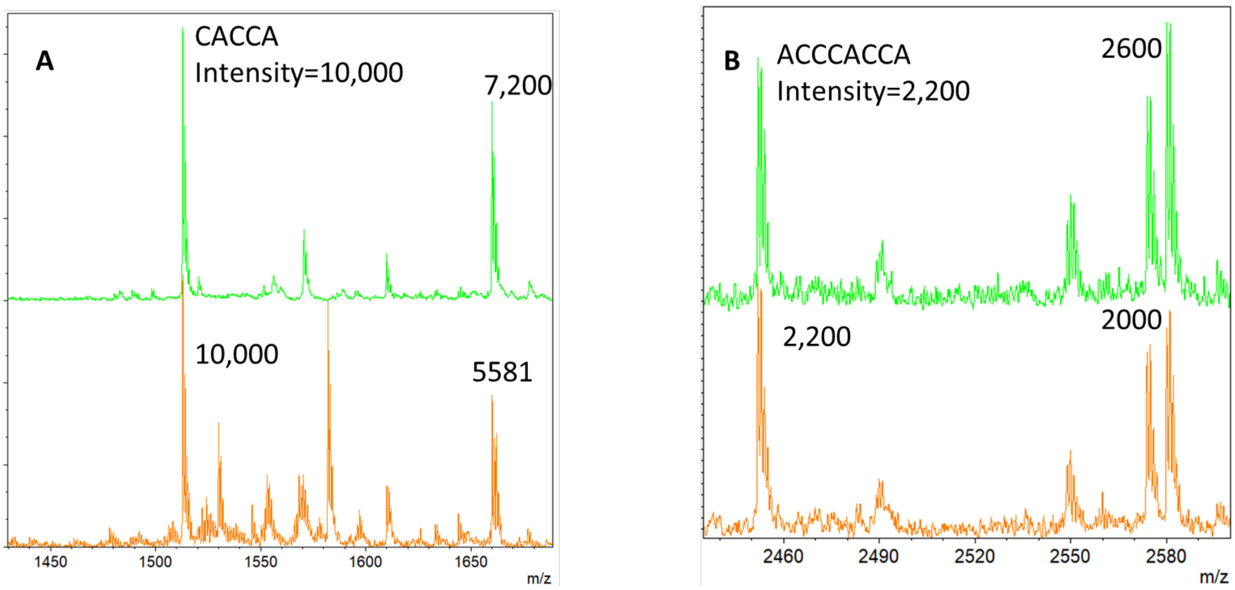


**Figure 4.** MALDI-TOF mass spectra of RNA-Phe (A) and RNA-Lys (B), respectively. The uncharged RNA peaks are normalized, and the relative charging efficiencies for D- (green traces) and L- (orange traces) are similar in both (A) and (B).

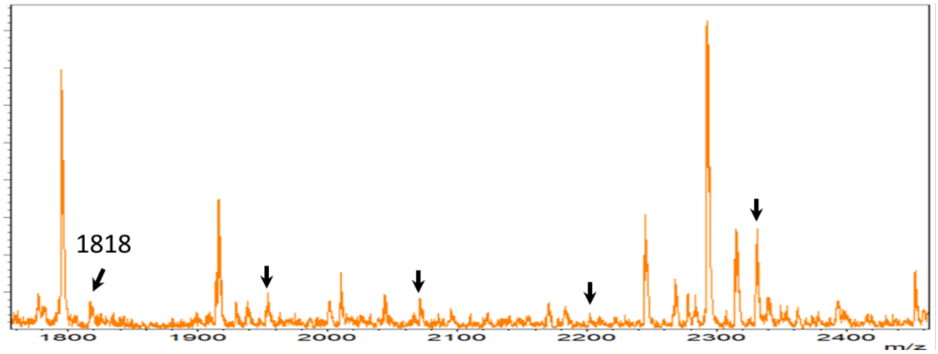


| | exp | cal |
|---|---|---|
| 1K | 1952 | 1946 |
| 2K | 2071 | 2074 |
| 3K | 2203 | 2203 |
| 4K | 2332 | 2331 |

**Figure S5.** Mass Spec relevant to Lysine oligos on CCACCA. For clarity, only incubation for L-lysine reaction is shown.

Table S1. Masses of common fragments in the main text.

| molecule | Mass |
|---|---|
| A (include PO3) | 329.2 |
| A-base | 135.1 |
| C (include PO3) | 305.2 |
| C-base | 111.1 |
| CACCACCA | 2452 |
| CCACCA | 1818 |
| CACCA | 1513 |
| ACCA | 1208 |
| + lysine | +128.2 |
| + Phenylalanine | +147.2 |

Experimental reagents and procedures:

1. T7 transcription.

All of the RNA samples (ptc1a/b) were synthesized using the HiScribe™ T7 High Yield RNA Synthesis Kit from New England Biolabs. After a 2-hour incubation, the reaction mixture volume was increased to 50 μL and treated with Dnase I for 15 minutes at 37°C. The mixture was then subjected to phenol extraction, and the aqueous phase was purified using the Monarch RNA Cleanup Kit (New England Biolabs). The quality of the RNA was assessed using a 3% agarose gel electrophoresis, as previously reported. For smFRET experiments, the RNA ptc1b is biotinylated at its 5'-end, and both ptc1a and 1b RNAs are labeled with Cy3 and/or Cy5 at the 3'-end with the commerical kit from Vector laboratory as described before.(1)

2. Preparation of tRNA synthetases.

The recombinant His-tagged $tRNA^{lys\ or\ Phe}$ synthetase was expressed in BL21(DE3) cells and purified on HisTrap 5ml on FPLC instrument (GE healthcare). Briefly, a single colony from a ampicilin agar plate is innoculated overnight in 100 ml of LB broth. The next day, this inoculation solution is diluted into 3 L of LB broth and cultured for approximately 3 hours until the absorbance (A590) reaches 0.8-1.2. Then, 1 mM IPTG is added to continue the culture for another 2-3 hours to induce the expression of the recombinant protein. After harvesting the cells by centrifugation and resuspending them in HisTrap buffer A along with a protease inhibitor cocktail, a pinch of lysozyme is added to the cell solution and the solution is frozen overnight at -30°C. The next day, the solution is thawed at 4°C for one hour and 1 μl of Dnase I is added and incubated on ice for another hour. The solution is then sonicated to break the cells and centrifuged twice, first at 6,000 xg for 30 minutes and then at 15,000 xg for 30 minutes, to remove debris.

The supernatant solution is passed through a 0.45 syring filter and loaded on a 5ml Histrap column ran by GE FPLC instrument. The recombinant protein is eluted from the HisTrap column using gradient elution of buffers A and B, in which B contains 1 M imidazole in addition to the components of buffer A. The His-tagged protein is eluted using a 200 mM imidazole solution. The identity of the protein is confirmed by its correct migration on a gel and by the large amount of protein present in the cell lysate, indicating over-expression. The FPLC fractions containing the protein are then pooled and concentrated, and the buffer is exchanged using Centricon filters. The final protein storage buffer is 20 mM tris-Hcl, pH 7.8; 10 mM $MgCl_2$; 4 mM BME; 0.5 mM EDTA; 400 mM KCl. The proper amount of synthetase is determined by analytical charging of total tRNAs.

3. Charging of L-, D- lysine/Phenylalanine to tRNA.

The charging mixture contains 100 mM Tris (pH 7.8), 4 mM ATP, 10 mM $MgCl_2$, 0.5 mM EDTA, 2 microM pure synthetase (or 10% volume of total synthetase), 7 mM BME, 20 microM tRNA, 100 microM of AA. The solution is incubated at 37°C for 15 min. Next, 1/10 volume of 20% KAC (pH 5) is added, followed by the addition of 1 volume of ultra-pure phenol (pH 4.5, from Thermo Fisher). The tRNA is then precipitated by adding 2 volumes of absolute ethanol and recovering the tRNA by centrifugation at 20,000 xg for 10 minutes. The charged tRNAs are then purified using a Monarch RNA cleanup kit from New England Biolabs.

4. Rnase T1 digestion. The charged and purified tRNAs are resuspended in water to a concentration of approximately 75A/ml. RNase T1 (50u/microliter) is then added to a 1/10th volume, and the digestion is incubated 37 °C for 15 minutes. Subsequently, the enzyme is removed by phenol extraction. The digested RNA -AA chimeras are purified using 20% PAGE mini-gel without urea (300V/30 minutes). The appropriate gel bands are visionilzed via UV radiation against a fluorescent silica TLC plate, and incised and eluted with water at 4°C for 1-2 hours. The resulting solution contains the purifed RNA-AA chimeras.

5. RNA-AA labeling with NHS dyes and gel observation.

The RNA-AA chimeras are mixed with 1 mM NHS dye in water and incubated at 4°C overnight.(1) The free dyes are then removed by passing the mixture through a Zeba desalting column twice. The Zeba desalting step helps to remove any unbound dye and ensure the purity of the labeled RNA-AA chimeras.

The labeled RNA-AA chimeras are loaded onto a 20% PAGE gel, with approximately 1-5 pmol being loaded based on the dye measurement. The gel is run for 30 minutes at 300 volts with a blue-ice coolant in the tank to separate the components based on size. The bands are immediately visualized using a BioRad ChemiDoc imager.

1. Xu, D. and Wang, Y. (2021) smFRET study of rRNA dimerization at the peptidyl transfer center. *Biophys. Chem.*, **277**, 106657.
